# Dynamic evolution of infarct volumes at MRI in ischemic stroke due to large vessel occlusion

**DOI:** 10.1101/2023.12.12.571221

**Authors:** Fanny Munsch, David Planes, Hikaru Fukutomi, Gaultier Marnat, Thomas Courret, Emilien Micard, Bailiang Chen, Pierre Seners, Johanna Dubos, Vincent Planche, Pierrick Coupé, Vincent Dousset, Bertrand Lapergue, Jean-Marc Olivot, Igor Sibon, Michel Thiebaut de Schotten, Thomas Tourdias, the FRAME and ETIS investigators

**Affiliations:** Univ. Bordeaux, Institut de Bio-imagerie IBIO, F-33000 Bordeaux, France; CHU de Bordeaux, Neuroimagerie diagnostique et thérapeutique, F-33000 Bordeaux, France; Kansai Electric Power Hospital, Osaka, Japan; CHRU Nancy, Inserm CIC-IT U1433, F-54511 Nancy, France; Institut de Psychiatrie et Neurosciences de Paris (IPNP), INSERM U1266, F-75014 Paris, France; Hopital Fondation Rothschild, departement de neurologie, F-75012 Paris, France; Univ. Bordeaux, CNRS, UMR 5293, Institut des Maladies Neurodégénératives, F-33000 Bordeaux, France; Univ. Bordeaux, Bordeaux INP, LABRI, CNRS, UMR5800, F-33400 Talence, France; Univ. Bordeaux, INSERM, Neurocentre Magendie, U1215, F-3300 Bordeaux, France; Hôpital FOCH, Service de Neurologie et Unité de Neuro Vasculaire, F-92151 Suresnes, France; CHU de Toulouse, unité neurovasculaire, F-31000 Toulouse, France; CHU de Bordeaux, Unité neurovasculaire, F-33000 Bordeaux, France; Univ. Bordeaux, CNRS, UMR-5293, F-33000 Bordeaux, France; Brain Connectivity and Behaviour Laboratory, Paris, France

## Abstract

**Background and Objectives:** The typical infarct volume trajectories in stroke patients, categorized as slow or fast progressors, remain largely unknown. This study aimed to reveal the characteristic spatiotemporal evolutions of infarct volumes caused by large vessel occlusion (LVO) and show that such growth charts help anticipate clinical outcomes.

**Methods:** We conducted a secondary analysis from prospectively collected databases (FRAME, 2017– 2019; ETIS, 2015–2022). We selected acute MRI data from anterior LVO stroke patients with witnessed onset which were divided into training- and independent validation-datasets. In the training-dataset, using Gaussian mixture analysis, we classified the patients into three growth groups based on their rate of infarct growth (diffusion volume / time-to-imaging). Subsequently, we extrapolated pseudo-longitudinal models of infarct growth for each group and generated sequential frequency maps to highlight the spatial distribution of infarct growth. We used these charts to attribute a growth group to the independent patients from the validation-dataset. We compared their 3-month modified Rankin scale (mRS) with the predicted values based on a multivariable regression model from the training-dataset that used growth group as independent variable.

**Results:** We included 804 patients (median age, 73.0 years [IQR, 61.2-82.0 years]; 409 men). The training-dataset revealed non-supervised clustering into 11% (74/703) slow, 62% (437/703) intermediate, and 27% (192/703) fast progressors. Infarct volume evolutions were best fitted with a linear (r=0.809; *P*<.001), cubic (r=0.471; *P*<.001), and power (r=0.63; *P*<.001) functions for the slow, intermediate and fast progressors, respectively. Notably, the deep nuclei and insular cortex were rapidly affected in the intermediate and fast groups with further cortical involvement in the fast group. The variable “growth group” significantly predicted 3-month mRS (multivariate OR, 0.51; 95% CI: 0.37-0.72, *P*<.0001) in the training-dataset, yielding a mean AUC of 0.78 (95% CI: 0.66-0.88) in the independent validation-dataset.

**Conclusions:** We revealed spatiotemporal archetype dynamic evolutions following large vessel occlusion stroke according to three growth phenotypes called slow, intermediate and fast progressors, providing insight into anticipating clinical outcome. We expect this could help in designing neuroprotective trials aiming at modulating infarct growth prior EVT.

## Introduction

In the event of large vessel occlusion stroke (LVOS), a cascade of cellular changes arises rapidly (1). This cascade begins with reversible electrical dysfunction (*i.e.,* ischemic penumbra) and progresses to ion pumping and energy metabolism failures inducing growth of the irreversible infarct core if cerebral blood flow is not restored (1). The infarct growth rate (IGR) varies substantially from one patient to another (2), influenced by various factors - most notably the ability of leptomeningeal collateral circulation to maintain residual perfusion, thereby minimizing the expansion of the infarct core into the penumbra (3, 4). Endovascular thrombectomy (EVT) is the standard-of-care treatment to reopen anterior LVO and stop the ischemic cascade (5). The rate of infarct progression prior to EVT, which is now referred as slow - intermediate - or fast progressors (6, 7), represents the inter-individual tolerance to ischemia and strongly correlates with clinical outcome after EVT (8). It is therefore crucial to improve our understanding on the growth phenotype prior to EVT which will be one of the future therapeutic target to continue improving the patients. Indeed, the growth phenotype could guide adjunctive therapy (9). The future challenge will be to interfere with stroke progression prior EVT which will require cerebroprotectant trials that will necessarily have to tailor specific neuroprotective strategy to specific growth phenotypes.

However, a major limitation is that there are currently no standard criteria to define slow, intermediate, or fast progressors, leaving us reliant on somewhat arbitrary definitions and thresholds (8, 10–13). For example, fast progressors have been arbitrarily characterized by an infarct volume higher than 70mL within the first 6 hours (7, 13), or an IGR higher than 5 mL/h (12) or 10 mL/h (8) among other definitions. One reason is the absence of comprehensive understanding of the archetypical dynamic evolutions of infarct volumes for the different growth phenotypes. The major challenge is that revealing the natural development of the infarct is impossible with a standard longitudinal approach because the vast majority of patients are scanned a single time before attempting rapid recanalization whenever possible. Only few data reported two imaging sessions prior to recanalization (14). An alternative method would consist of inferring pseudo-longitudinal profiles from the concatenation of a large number of homogeneous cross-sectional data collected before recanalization akin to strategies used recently to map the typical evolution of brain volume across the lifespan (15, 16) or in neurodegenerative diseases (17, 18). Indeed, if we could objectively attribute stroke patients presenting the same clot location to a specific growth phenotype, then, we could reconstruct stereotypical growth curves from such homogeneous cluster of patients explored at different time points after the onset of symptoms.

Based on these considerations, we aimed to identify slow, intermediate and fast progressors following LVOS using non supervised clustering approach in order to reveal the characteristic spatiotemporal evolutions of infarct volumes according to such phenotypes of tolerance to ischemia. Additionally, we seek to provide the first evidence supporting the role of these new charts in anticipating patients’ outcomes.

## Methods

### Standard Protocol Approvals, Registrations, and Patient Consents

We conducted a secondary analysis using two prospective studies approved by the institutional ethics committee, the primary objectives of which were unrelated to the present article. These are the "French acute multimodal imaging to select patients for mechanical thrombectomy" (FRAME; NCT03045146) and the "Endovascular treatment in Ischemic stroke" registry (ETIS; NCT03776877). Written informed consent was obtained from all patients or their legal representatives. Our analysis called DEVOL (“Dynamic EVOLution of stroke”) was reported following the Strengthening the Reporting of Observational Studies in Epidemiology criteria for observational studies (19).

### Patients

FRAME prospectively recruited consecutive participants in 2 comprehensive stroke centers in France from January 2017 to February 2019 who presented with LVOS and underwent acute imaging including perfusion-weighted imaging (PWI) on arrival, and were subsequently treated with EVT (20). Analysis of the images from FRAME patients has already been reported but focusing on penumbra (20, 21) while we explored the IGR in the present paper.

ETIS is an ongoing, multicenter, prospective, real-life observational study evaluating LVOS treated with EVT in 21 comprehensive stroke centers in France. Images are now sent by the investigative centers and stored centrally (ETIS-image subproject). There is a manual step of verification of the images and proper identification before their upload in a dedicated environment (ArchiMed). We considered the images available in ArchiMed for patients explored between January 2015 and January 2022. Details regarding data collection and materials have been previously published (22). Analysis from the images of ETIS patients has not been reported yet.

Within FRAME and ETIS databases, we selected the patients with:

1. Anterior circulation LVO (intracranial internal carotid artery, M1 segment of the middle cerebral artery, tandem with proximal intracranial occlusions),
2. Witnessed stroke onset,
3. MRI as baseline imaging.

We excluded patients with stroke-on-awakening or unprecise onset, as well as those explored with computer tomography or whose MRI was of insufficient quality (**Fig-1**).

**Figure 1:**
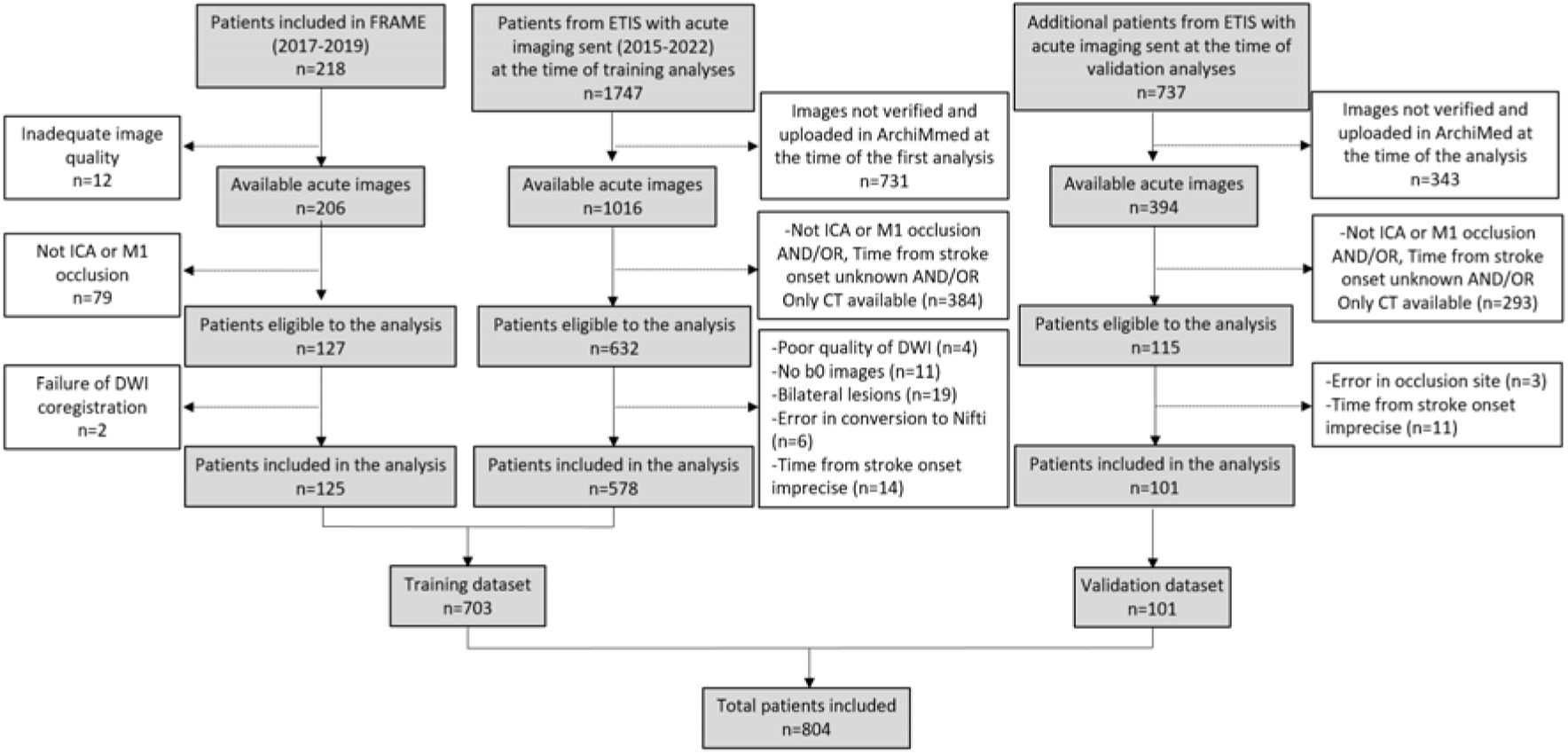
Flowchart. Inclusion and exclusion for patients from FRAME and ETIS.

We accessed a first set of available data of 703 patients from the FRAME and ETIS cohorts used as a training dataset to model temporal, spatial and clinical evolution. Then, later we got access to 101 additional new patients from the ETIS dataset to build an independent dataset for validation (**Fig-1**).

### Image analyses

All the images were analyzed centrally blinded to clinical data. The infarct core masks, used to measure infarct volume and calculate IGR in our study, were delineated from acute diffusion MRI using automatic detection from Olea Sphere (version SP34) and manual correction afterwards if needed. Then, we registered all the images and the associated core masks to the standard Montreal Neurologic Institute 152 (MNI152) template as described previously (21).

PWIs from FRAME had already been reported elsewhere (20, 21), and we processed the additional PWI available from ETIS with Olea Sphere (version SP34). The masks of time-to-maximum (Tmax) greater than 6 seconds and 10 seconds were automatically extracted, visually checked and normalized to the MNI152 template.

We derived collateral status from the hypoperfusion intensity ratio (HIR) that is the percentage of the perfusion lesion with severely delayed contrast arrival times according to the formula: HIR = volume of Tmax>10s / Tmax>6s. HIR was repeatedly identified as a reliable proxy of good collaterals defined by angiography when HIR<0.4 (23–25). When PWI was unavailable, we used data from the conventional angiography performed before EVT (if adequate with sufficiently late venous phases), and considered grades 3 and 4 from the ASITN/SIR classification as good collateral status (26).

We also computed the volume of perfusion (Tmax >6s) / volume of the core infarct (diffusion) mismatch to estimate the tissue at risk.

### Temporal evolution of infarct volumes

We defined IGR as the ratio of infarct core volume on acute diffusion MRI to the time between stroke onset and imaging. We explored the distribution of IGR within the training dataset and we used an unsupervised clustering approach to identify subtypes of stroke progressors among the entire dataset. We used a Gaussian mixture approach (27), constraining the model to 3 clusters according to the literature (7), later referred to as slow, intermediate, and fast progressors. We explored the different clinical and imaging characteristics between such defined growth groups with Kruskal Wallis rank sum tests.

Then, we extrapolated pseudo-longitudinal core volume models across time for each group. The rationale behind this approach is that the patients from the same growth phenotype and with the same type of proximal occlusion (all are ICA or M1) should share similar evolution and could be used to extrapolate volumetric trajectories. We have already showed that such concatenation of several cross-sectional data is able to infer the same dynamic as “truly” longitudinal data (17). We first used the non-parametric LOESS approach (locally estimated scatterplot smoothing) consisting of fitting models to localized subsets of the data point by point (28) to represent smooth curves. We also tested ten models from the simplest to the most complex (linear, logarithmic, inverse, quadratic, cubic, power, S, growth, exponential, logistic) and kept the one with the highest r value.

### Spatial evolution of infarct volumes

We computed 3D frequency maps for 5 consecutive bins of 1h in each growth group using the core masks normalized in MNI152 after flipping all the infarcts to the left hemisphere to increase the statistical power. These maps represent the number of time each voxel was impacted by an infarct. We quantified the areas affected through frequency map projections into the Harvard-Oxford cortical and subcortical structural atlas (29). We tested the different involvement of the atlas regions according to time, group, and time * group using 2-way MANOVA with Wilks’ Lambda tests and correction for multiple comparisons.

The previous approach provided the typical infarct growth location according to the growth phenotype but only for 5 steps of delay from stroke onset (of 1h each).

### Impact of growth group to anticipate the clinical outcome

In the training dataset, we used logistic regression with the growth group as the primary independent variable to predict clinical outcomes, defined as good for a 3-month Rankin scale (mRS) ≤2. We checked multicollinearity among variables with Spearman correlations (**Supplementary-Fig-1**). We quantified the model performance with the area under the receiver operating characteristic curve (AUC) with bootstrap technique to estimate the 95% CI. Then, we applied this prognosis model to the patients from the independent dataset for external validation. We projected the core volumes of these new patients onto the longitudinal profiles created above to identify their growth groups from the smallest Euclidean distance to one of the three fitting curves. We compared the probability of these new patients having good outcome with the actual mRS and re-computed AUC and 95% CI. Analyses were implemented on R (version 4.2.2) and SPSS (version 29.0.1.0).

## Data availability

Data could be made available upon request to the principal investigators.

## Results

### Patient characteristics

Among the 218 participants enrolled in FRAME, we excluded 93 from this analysis for reasons summarized in the flowchart (**Fig-1**). Regarding ETIS, from January 2015, to January 2022, we counted 1747 patients with acute imaging. The images already verified, quality controlled and uploaded in the storage system (ArchiMed) were available for 1016 patients. Finally, after application of the inclusion criteria, 578 patients with acute imaging were included and added to the FRAME patients to create the training dataset of 703 patients (**Fig-1**). At the time of conducting the validation analyses (09/2022), an additional set of 101 patients was available from ETIS (**Fig-1**) and used as an external validation dataset.

The total of 804 patients consisted of 409 (50.9%) men, with a median age of 73.0 years (IQR, 61.2-82.0 years) and a median NIHSS score of 17.0 (IQR, 11.0-21.0). Most (n=746, 94.1%) were treated with EVT after a median delay of 4.2 hours (IQR, 3.1-5.5). Other patient characteristics are shown in Table 1. The training dataset and the validation dataset showed close characteristics (**Table-1**).

**Table 1:**
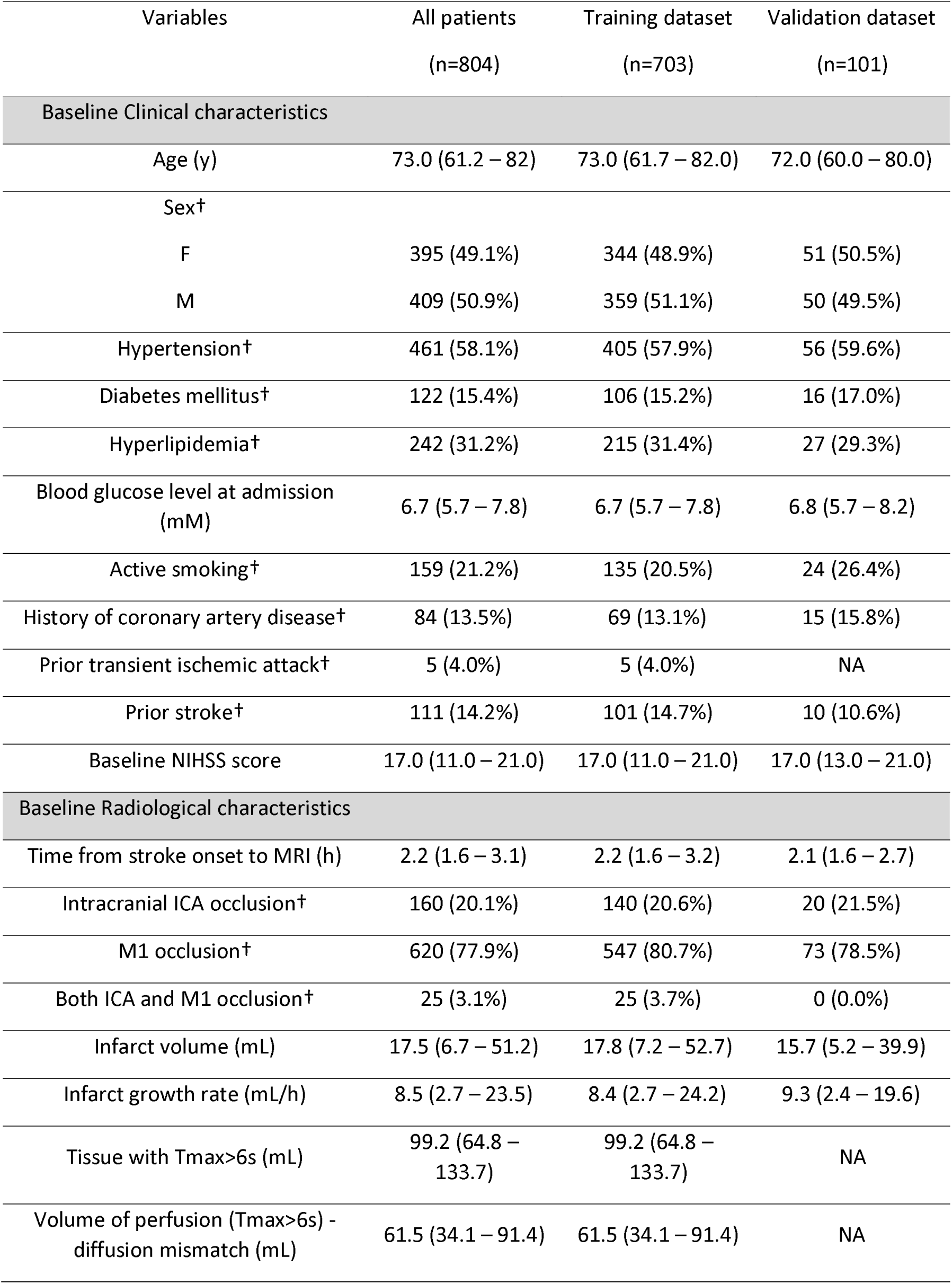
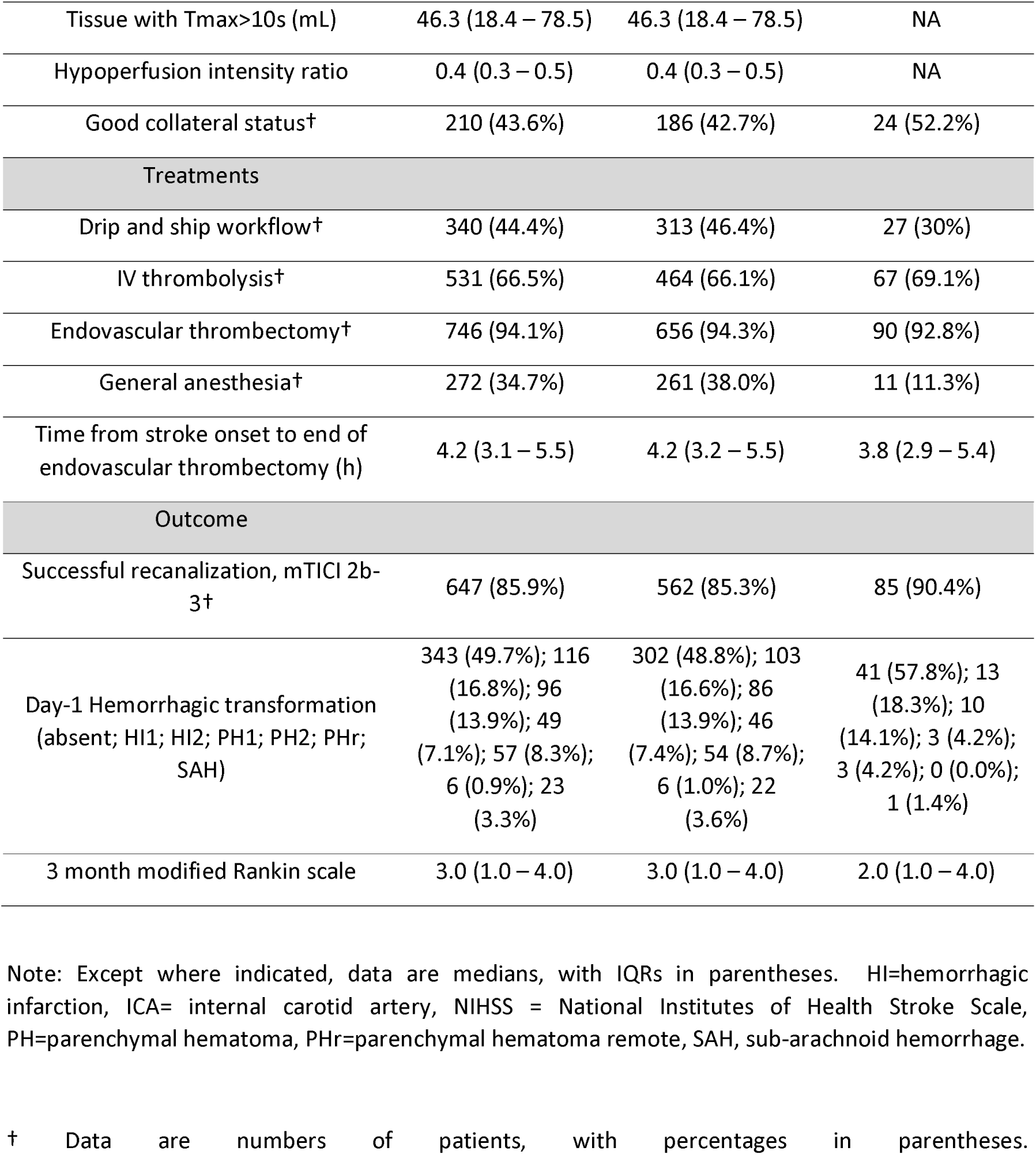
Characteristics of patients.

### Temporal evolution of infarct volumes

Within the training dataset, the median volume of infarct core before recanalization was 17.8 mL (IQR, 7.2-52.7) as observed on diffusion MRI collected after a median delay of 2.2 hours (IQR, 1.6-3.2, *Table-1*). This translated into a median calculated IGR of 8.4 mL/h (IQR, 2.7-24.2) that showed a non-normal left-skewed distribution (**Fig-2A**). After applying natural log-transformation, we got a more centered distribution, but the IGR spread over a wide range of values, pointing toward different sub-populations within this sample. Under the assumptions of three sub-populations, later called slow, intermediate, and fast progressors, we identified a mixture of three Gaussians within the overall sample (**Fig-2B**) that provided the likelihood for each patient to belong to a given group (**Fig-2C**). This non-supervised approach clustered groups representing respectively 11% (74/703), 62% (437/703), and 27% (192/703) of the total sample (**Table-2**).

**Figure 2:**
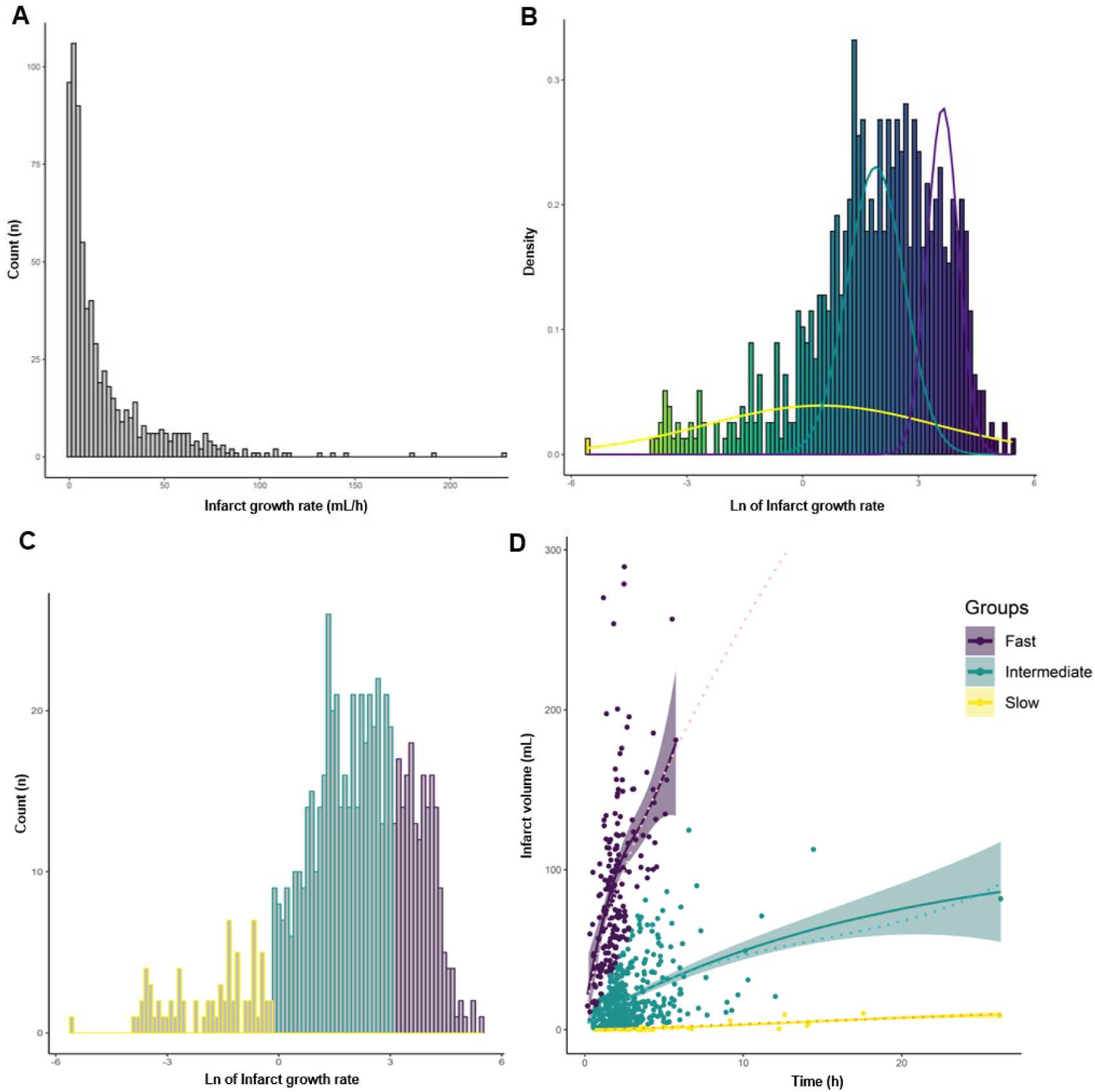
Distribution of infarct growth rate and temporal evolution of infract volumes. The histogram of infarct growth rate in the training dataset showed a non-normal left-skewed distribution **(A)**. After natural logarithmic transformation, the distribution of infarct growth rate was modeled as a mixture of three Gaussian shape subpopulations **(B)** which allowed the attribution of each patient to a slow, intermediate or fast progression profile **(C).** Panel **D** shows the distribution of infarct volumes according to the time from stroke onset for the training dataset. The dynamic evolution of infarct volumes has been modeled through concatenation of the cross-sectional data for the slow, intermediate and fast progressors. Solid lines show the locally estimated scatterplot smoothing and the associated confidence intervals while dashed lines show the best parametric fits corresponding to a linear function for the slow progressors (yellow), a cubic function for the intermediate progressors (cyan) and a power function for the fast progressors (purple).

**Table 2:**
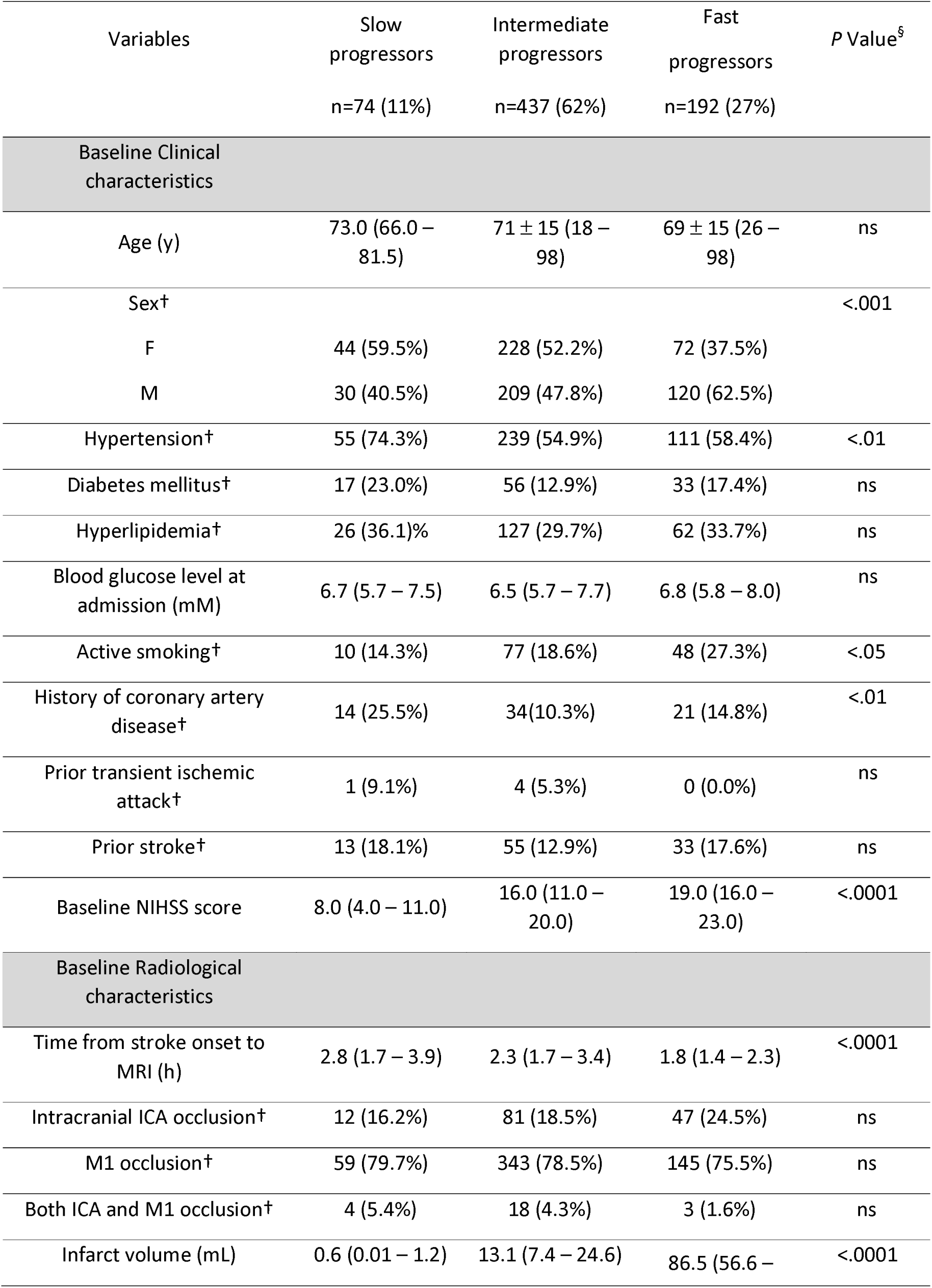
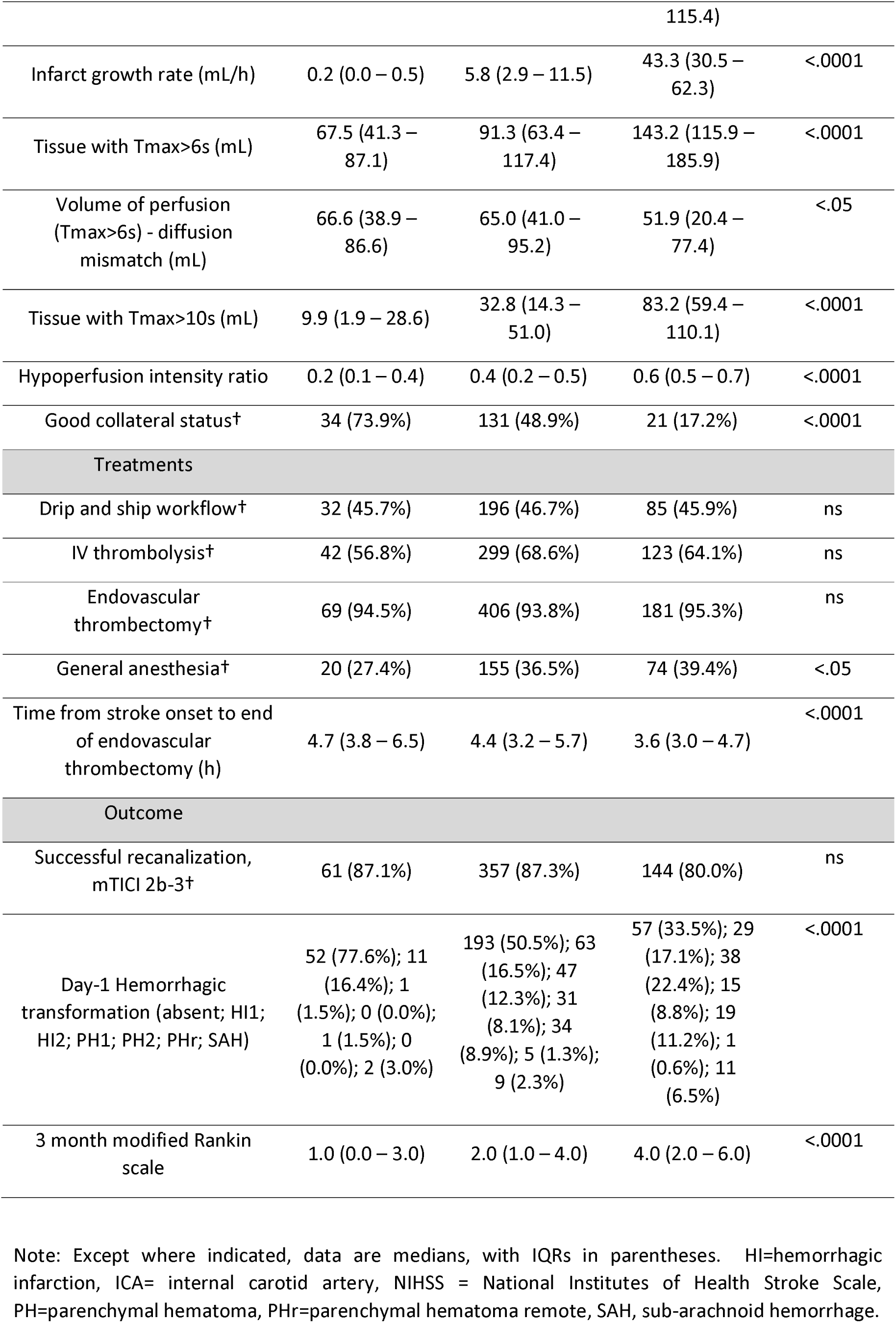
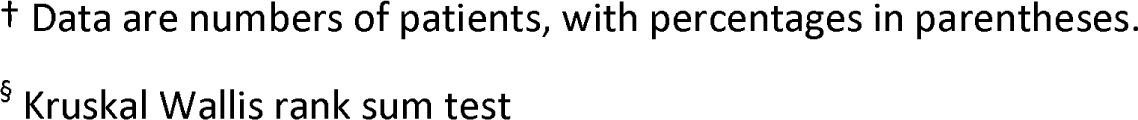
Characteristics of patients according to the growth group in the training dataset.

These growth groups showed significantly different IGR (median 0.2 mL/h [IQR, 0.0-0.5 mL/h] vs. 5.8 mL/h [IQR, 2.9-11.5 mL/h] vs. 43.3 mL/h [IQR 30.5-62.3 mL/h]; *P*<.0001) and different initial NIHSS severity (median 8.0 [IQR, 4.0-11.0] vs. 16.0 [IQR, 11.0-20.0] vs. 19.0 [IQR, 16.0-23.0]; *P*<.0001). A slower growth group was also significantly associated with a higher volume of mismatch and a significantly better collateral status (73.9% of good collaterals vs. 48.9% vs. 17.2 %; *P*<.0001). These patients all showed the same type of anterior LVO and were treated similarly with EVT combined with IV thrombolysis whenever eligible with a non-different rate of successful recanalization. However, the rate of high-grade hemorrhagic transformation was significantly higher from slow to intermediate to fast progressors, and the 3-month functional outcome worsened along the groups (median 3-month mRS of 1.0 [IQR, 0.0-3.0] vs. 2.0 [IQR, 1.0-4.0] vs. 4.0 [IQR, 2.0-6.0]; *P*<.0001). Because each of these 3 groups can be considered as a homogeneous cluster of patients explored at different time points after the onset of symptoms, we could infer the pseudo-longitudinal evolutions of core volumes by fitting the cross-sectional data of each group. LOESS fitting provided the estimated virtual pictures of the expected time-course evolution before recanalization according to the growth group (**Fig-2D**). As LOESS cannot produce a function that is easily represented by a mathematical formula to apply to new patients’ data, we also tested parametric models. The evolution of the slow progressors was best modeled with a low slope linear fitting (r=0.809; *P*<.001). The evolution of the intermediate progressors was the closest to a cubic function (r=0.471; *P*<.001), while the logarithmic fit was not far (r=0.452; *P*<.001). The evolution of the fast progressors was modeled with a power function (r=0.63; *P*<.001) with values close to an exponential fit (r=0.558; *P*<.001; **Fig-2D, dashed lines**).

### Spatial evolution of infarct volumes

We generated 3D frequency maps from the coregistered core masks to translate the above-defined time course evolutions into archetype infarct locations according to the growth group. The slow progressors showed no stereotypical infarct pattern but instead small and unpredictable lesions within the internal carotid artery territory (**Fig-3A**). By contrast, the intermediate and fast progressors showed stereotypical patterns (**Fig-3B and C**) whose global time courses were significantly different (time x group interaction; p<0.001; MANCOVA). In both groups, the pallidum, the putamen, the caudate, the insular cortex and the frontal operculum cortex were non-differentially affected from the first hours (time x group interaction; *P-*values close to 1; **Fig-4**). But the fast progressors were characterized by a quicker and more systematic involvement of the cortical parcels of the internal carotid artery territory, including clinically relevant regions such as the pre- and post-central gyrus, the angular, supramarginal, and the posterior division of the superior temporal gyrus. **Fig-4** summarizes the mean proportions of each cortical area affected during time.

**Figure 3:**
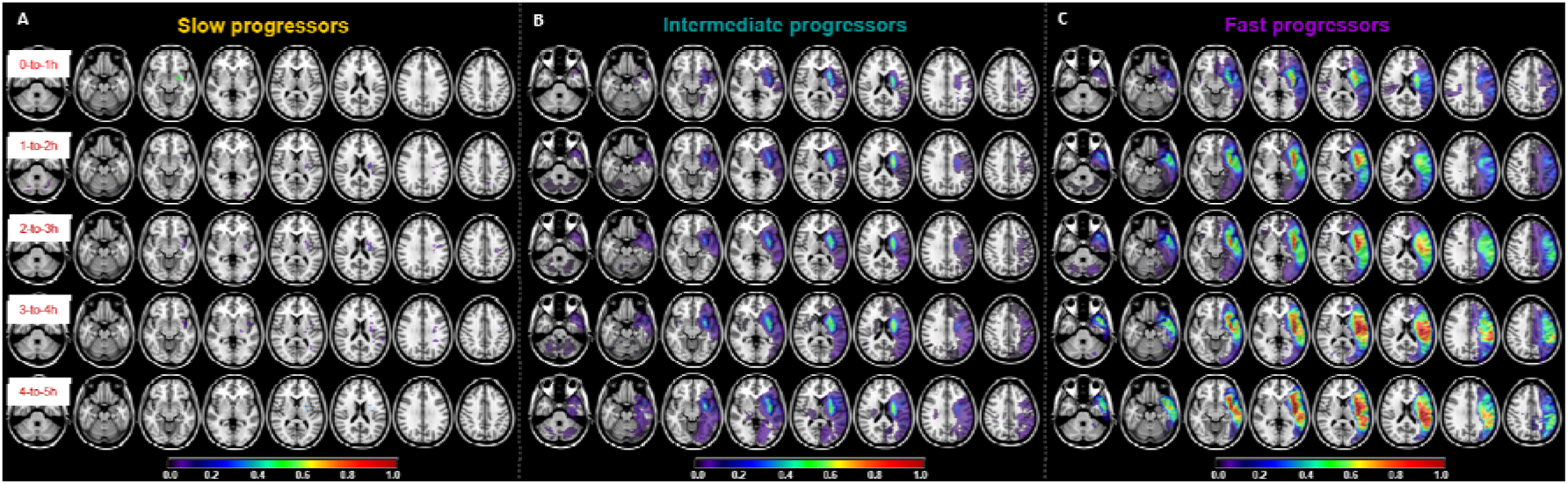
Spatial evolution of infarct volumes. Prevalence maps from the training dataset show the frequency of infarct locations for the slow **(A),** intermediate **(B)** and fast progressors **(C)** according to the time from stroke onset divided in 1 hour bin. Color coding indicates the ratio of overlapping lesions out of the total number of infarcts in each group and each bin. All the infarct cores have been flipped to the left size for this analysis. The maps are not shown above 5 hours because of the smaller sample size.

**Figure 4:**
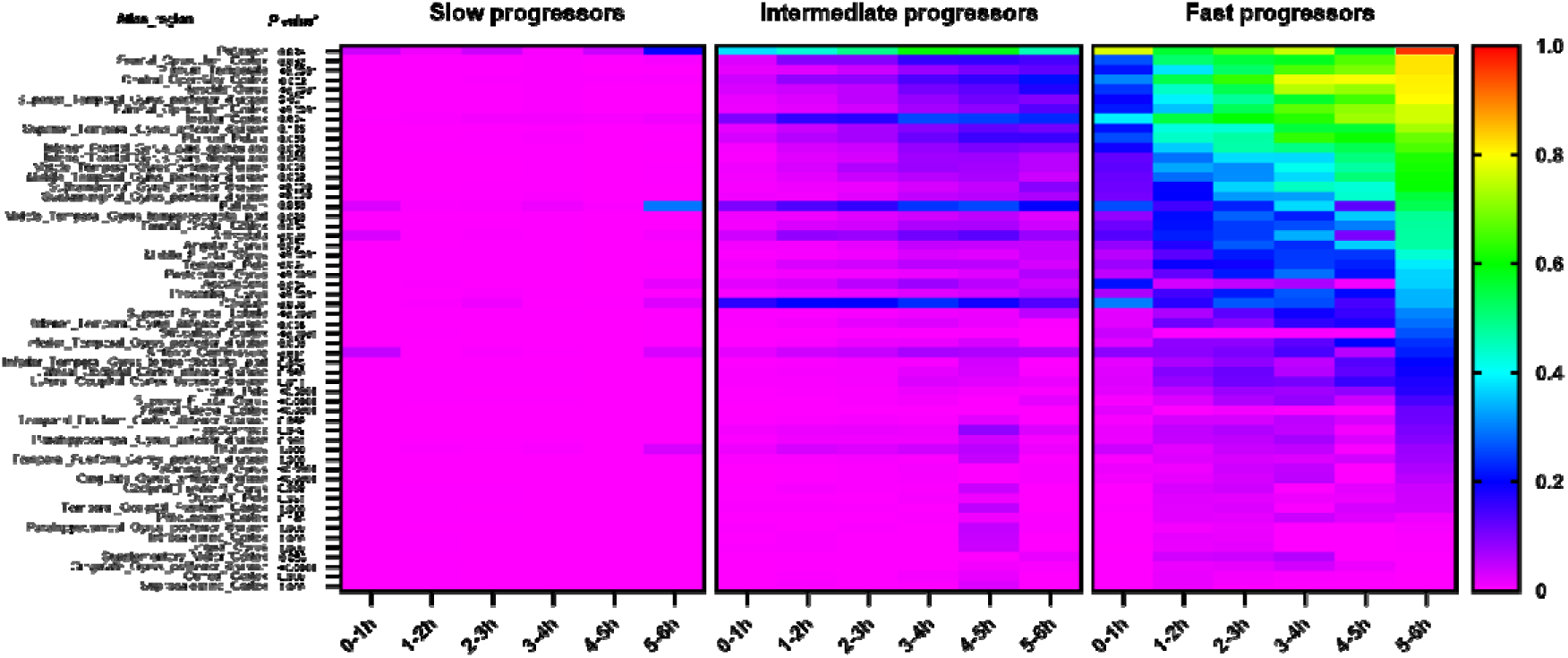
Heat maps of infarct location. The 60 cortical and subcortical parcels from the Harvard-Oxford atlas (29) are listed on the left. The heat maps show the mean proportions (color-coded from 0 to 1) of each parcel that are affected by the infarcts with time passing and according to the growth group in the training dataset. For each parcel p-values indicate the time * group interaction from the 2-way MANOVA analysis.

### Impact of growth group to anticipate the clinical outcome

We tested whether the above-defined dynamic evolutions of infarct volume could help anticipating new patients’ clinical outcome. Within the training dataset, we found that the growth group was a significant and independent predictor of favorable functional outcome at 3 months (OR, 0.51; 95% CI: 0.37-0.72; *P*<.0001) together with age (OR, 0.96; 95% CI: 0.95-0.97; *P* <.0001) and initial NIHSS severity (OR, 0.91; 95% CI: 0.88-0.94; *P<*.0001). Overall, we found mean discrimination in terms of AUC of 0.75 [95%CI: 0.71, 0.79]. See **Table 3** and **Supplementary-Fig-2-A**.

**Table 3:** Logistic regression model for prediction of good functional outcome (mRS≥2) on the training dataset (n=703)

| Variable | Univariable analysis |  | Multivariable analysis |  |
| --- | --- | --- | --- | --- |
|  | Odds ratio | *Corrected*<br>*P* value | Odds ratio | *P* value |
| Age (+1 y) | 0.96 (0.95 – 0.97) | <.0001 | 0.96 (0.95 – 0.97) | <.0001 |
| Sex (F) | 0.78 (0.57 – 1.06) | ns |  |  |
| Hypertension (yes) | 0.68 (0.49 – 0.93) | ns |  |  |
| Diabetes mellitus (yes) | 0.58 (0.37 – 0.90) | ns |  |  |
| Hyperlipidemia (yes) | 0.89 (0.64 – 1.26) | ns |  |  |
| Blood glucose level at admission (+ 1 mM) | 0.87 (0.80 – 0.95) | ns |  |  |
| Active smoking (yes) | 1.70 (1.13 – 2.56) | ns |  |  |
| History of coronary artery disease (yes) | 0.87 (0.50 – 1.48) | ns |  |  |
| Prior transient ischemic attack (yes) | 1.43 (0.23 – 11.12) | ns |  |  |
| Prior stroke (yes) | 0.51 (0.32 – 0.81) | ns |  |  |
| Baseline NIHSS score (+ 1 point) | 0.88 (0.86 – 0.91) | <.0001 | 0.91 (0.88 – 0.94) | <.0001 |
| Time from stroke onset to MRI (h) | 1.07 (0.10 – 1.15) | ns |  |  |
| Intracranial ICA occlusion (yes) | 0.80 (0.58 – 1.09) | ns |  |  |
| M1 occlusion (yes) | 1.49 (1.04 – 2.15) | ns |  |  |
| Infarct volume (+ 1 mL) | 0.98 (0.98 – 0.99) | <.0001 |  |  |
| Infarct growth rate (+ 1 mL/h) | 0.97 (0.97 – 0.98) | <.0001 |  |  |
| Growth group (slow, intermediate, fast) | 0.41 (0.31 – 0.54) | <.0001 | 0.51 (0.37 – 0.72) | <.0001 |
| Tissue with Tmax>6s (+ 1 mL) | 0.99 (0.99 – 1.00) | <.05 |  |  |
| Volume of perfusion (Tmax>6s) - diffusion mismatch (+1 mL) | 1.00 (0.99 – 1.00) | ns |  |  |
| Tissue with Tmax>10s (+ 1 mL) | 0.99 (0.98 – 1.00) | <.05 |  |  |
| Hypoperfusion intensity ratio | 0.22 (0.06 – 0.75) | ns |  |  |
| Good collateral status (yes) | 1.74 (1.17 – 2.60) | ns |  |  |
| Drip and ship workflow (yes) | 0.76 (0.55 – 1.04) | ns |  |  |
| IV thrombolysis (yes) | 1.93 (1.38 – 2.70) | <.01 |  |  |
| Endovascular thrombectomy (yes) | 0.92 (0.45 – 1.89) | ns |  |  |
| General anesthesia (yes) | 0.10 (0.74 – 1.34) | ns |  |  |
| Time from stroke onset to end of endovascular thrombectomy (+ 1h) | 0.99 (0.93 – 1.05) | ns |  |  |
| Successful recanalization, mTICI 2b-3 (yes) | 3.05 (1.87 – 5.16) | <.001 |  |  |
| Day-1 Hemorrhagic transformation (no) | 3.66 (2.60 – 5.20) | <.0001 |  |  |

Note: Data in parentheses are 95% CIs. In univariable analysis, p-values have been corrected for multiple comparisons (Bonferroni corrected for 29 comparisons). In the multivariable analysis, we kept only the variables that could be available at the patient admission which is not the case for the variables related to the type of treatment or its outcome (IV thrombolysis, endovascular thrombectomy, general anesthesia, time from onset to the end of endovascular thrombectomy, successful recanalization, and hemorrhagic transformation). There was a significant collinearity between the variables “growth group”, “infarct volume”, “infarct growth” and “tissue with Tmax>6s” which was expected. We favored the growth group in the multivariable analysis as it is the primary interest of this analysis.

The validation dataset included 87 patients who had a mRS at 3 months from the 101 available. Within this sample that was not used to build the above-mentioned predictive model, we could project each patient onto the stereotypical time course evolutions defined in **Fig-2D**, which allowed us to attribute a growth group to each of these new patients according to the smallest Euclidean distance with one of the three infarct growth fits. We classified 34% (30/87) of patients as slow progressors, 51% (44/87) as intermediate, and 15% (13/87) as fast progressors. Using these groups and the previously defined predictive model, we predicted mRS at 3 months with an AUC of 0.78 [95%CI: 0.66, 0.88]. The calibration plot showed fairly good correspondence between the predicted probability of good outcome and the observed outcome (**Supplementary-Fig-2-B and C**).

## Discussion

The individual tolerance to ischemia following large vessel occlusion varies significantly among patients, leading to the recent terminology of slow or fast progressors. But no consensual definitions exist for such phenotypes whose dynamic courses needed to be discovered. Here, we employed, for the first time to our best knowledge, a data-driven non-supervised approach to delineate distinct categories of stroke progressors: slow, intermediate, and fast, and we unveiled the characteristic dynamic evolutions of these groups in both time (with volume along time charts) and space (with 3D frequency maps), while shedding light on their clinical relevance. Indeed, using these charts, new patients from an independent validation dataset could be classified in a progressor category, and their probabilities of good outcome could be predicted. In the future, this may help decide therapeutic strategies and may guide patient selection criteria for future neuroprotective trials.

The ambiguity surrounding the definition of slow and fast progressors has been a notable challenge to transfer this otherwise simple concept into clinical practice. For instance, "fast" has been defined as a core volume >70ml within the first 6 hours and "slow" <30ml between 6h and 24h (7, 13), but also according to evolution of the ASPECT score (9, 10, 30), to dichotomy of IGR based on the median value within a sample (12, 31), to tertile (11), or to a value that best correlates with the clinical outcome (8). To address this issue, we opted for a data-driven, non-supervised approach using a soft clustering mixture model. This methodology avoids reliance on arbitrary thresholds and instead identifies groups that bear similarities to those described in previous literature. Notably, our results align with existing estimates, suggesting that approximately 20%-to-30% of patients with LVOS may fall into a fast-to-ultra-fast progressor category (6), a proportion consistent with our 27% fast progressors. Our intermediate group could resemble those who were previously categorized as "slow" progressors, while we identified a smaller subset (∼11%) as "slow" progressors who exhibit minimal growth despite proximal occlusion.

One significant contribution of our study lies in utilizing over 700 core volumes, all coregistered within the same standard space, to infer longitudinal profiles of untreated stroke progression, drawing inspiration from methodologies applied in other domains (15–18). In contrast to previous efforts that employed linear fitting (32), which simplifies the complex biological processes at play, we employed smooth curve fitting. This approach revealed the non-linear nature of infarct growth. Our observations were consistent with serial MRI scans in animal stroke models, which indicated a logarithmic pattern for the natural evolution of infarct volumes (33, 34). In the case of fast progressors, results indicated that a power function provided a better fit, reflecting a steeper initial increase than the logarithmic pattern. Additionally, we noted that an exponential shape could describe the trajectory of fast progressors in the initial hours but failed to account for asymptomatic growth linked to the maximum volume of the vascular territory. Further investigations, such as trajectory-based clustering (35) might be beneficial when leveraging more extensive databases from international networks and data-sharing initiatives.

Our 3D frequency maps provided a spatial dimension to the temporal progression of stroke, offering insights into regions that exhibit varying vulnerability or tolerance to ischemia. We could revisit the known susceptibility of the deep nuclei and the insular cortex, which is in line with frail pial collateral support of these regions (36). Indeed, collateral status emerged as a key determinant of our growth profiles, a finding consistent with previous research (3, 4).

Furthermore, our growth group classification proved valuable in predicting functional outcomes. The predicted probability of good outcome of patients from the validation dataset were close to the observed one, with an AUC of 0.78 [95%CI: 0.66, 0.88]. It is, however, essential to acknowledge the inherent challenges in making acute-stage predictions with only pre-treatment information. Though, we believe the method to be sufficiently accurate to capture groups with homogeneous expected outcome which could help to maximize statistical power in future cerebroprotectant trials. Indeed, an effective strategy interfering with infarct growth prior EVT (such as during transfer to a comprehensive stroke center) will have a higher expected effect on fast progressors than on the other phenotypes which should translate into statistical demonstration of its efficacy with a smaller number of participants if only fast progressors are included versus unselected patients.

Several limitations warrant consideration. We focused on patients considered eligible for EVT, potentially excluding certain patients, particularly in the fast progressor group. Nonetheless, our inclusion of close to 30% of fast progressors surpassed the proportion within initial clinical trials that often involve favorable profiles (37). In that sense, the ETIS real-life prospective registry, representing the largest reported population from multicenter acquisitions, is a strength for translating our results to daily clinical practice. The utilization of MRI limited the generalizability of our findings to settings where CT is the primary imaging modality. However, this approach, which is the first line imaging in several French centers, together with inclusion of witnessed symptom onset only which was not systematic in prior works (8, 11), provided high accuracy for growth quantifications. We also acknowledge the need for collective efforts in the future to incorporate more data from extended time windows to generalize our IGR charts as most of our patients presented within 3 hours from the onset. We will also have to explore the impact of recanalization on spatiotemporal dynamics. Additionally, we based our external validation on the clinical outcome because we could not access a natural history follow-up imaging to check if the infarct growth followed the prediction. This could be tested in the future with subgroup of patients whose recanalization failed. Finally, PWI was not always available and therefore we could not compute HIR for all the patients and we also assessed collateral status through conventional angiography when PWI was not available. We acknowledge that pre-thrombectomy angiographies don’t necessarily include late series and that combining HIR and ASITN/SIR is suboptimal even though several papers showed that both metrics are correlated (23–25). But even though our collaterality metric is imperfect, the association of collateral status with the growth phenotypes was already reported (4) and was not the main objective of our work that rather focused on revealing unbiased growth charts for infract progression in time and space.

In conclusion, our study has unveiled infarct growth charts for stroke progression that offer a novel and practical approach for phenotype identification in stroke patients and outcome estimation, all without relying on advanced post-processing techniques. In the future, these growth charts could aid the design of neuroprotective trials and to personalize the treatment strategy.

## Data sharing

Data generated or analyzed during the study are available from the corresponding author by request.

## List of abbreviations

ASITN/SIR: American Society of Interventional and Therapeutic Neuroradiology/Society of Interventional Radiology
EVT: Endovascular thrombectomy
IGR: Infarct growth rate
LOESS: locally estimated scatterplot smoothing
LVO: Large vessel occlusion
LVOS: large vessel occlusion stroke
mRS: modified Rankin scale
PWI: perfusion weighted imaging

