## Supplementary material for "Dynamic evolution of infarct volumes at MRI in ischemic stroke due to large vessel occlusion"

**Supplementary Figure 1:** Spearman correlation coefficients for each combination of variables.

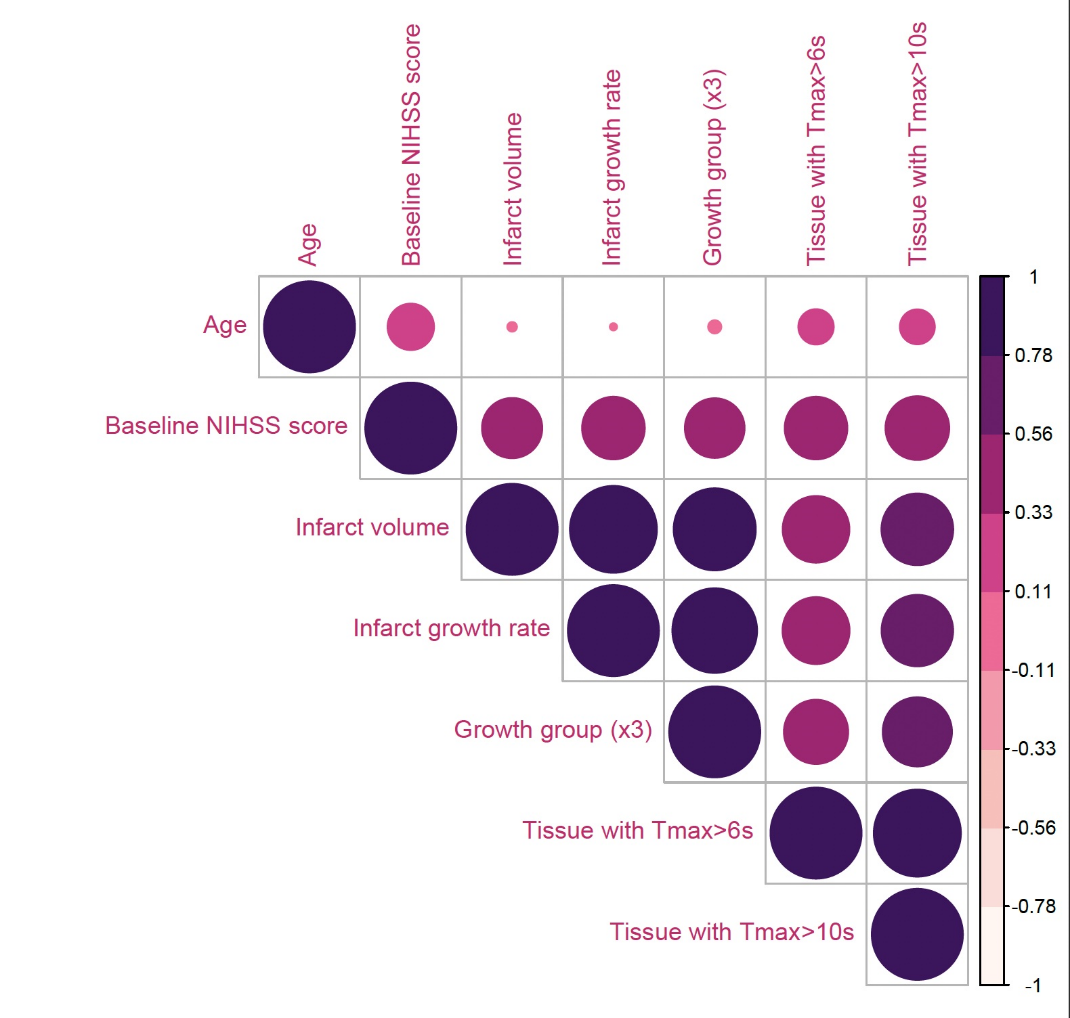

As expected, the growth group shows correlation with infarct volume, infarct growth rate but also with the severity of hypoperfusion which is why these variables were not included together in the final predictive model

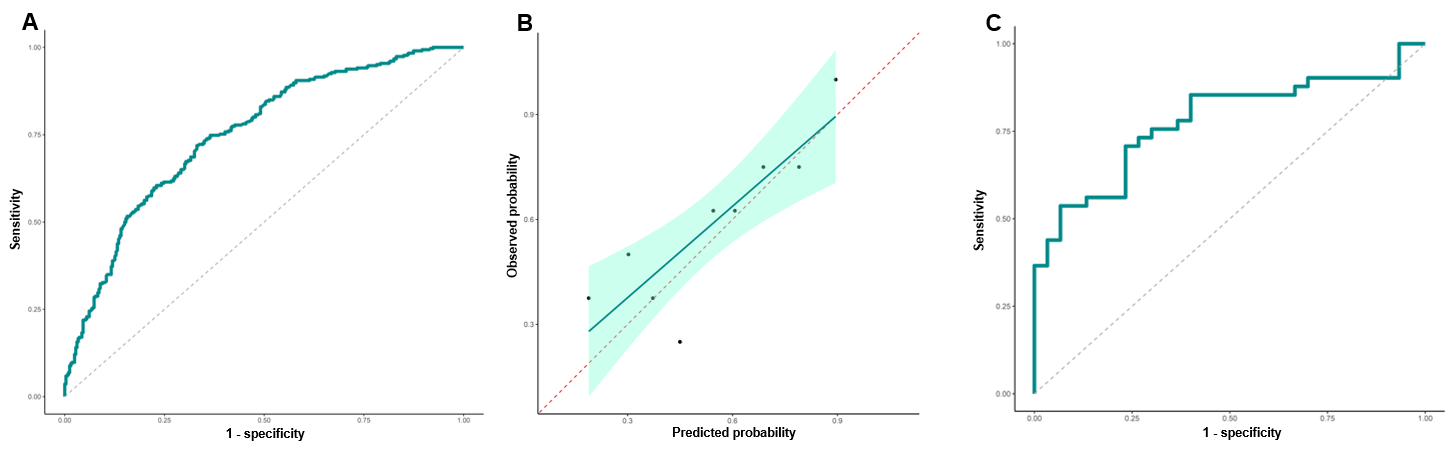
**Supplementary Figure 2:** Predictive performances of the model including the growth group

**(A)** Receiver operating characteristic curve of the predictive model for good outcome in the training dataset shows area under the curve of 0.75 [95%CI: 0.71, 0.79]. **(B)** Calibration plot shows good correspondence between the predicted probability of good outcome and the observed outcome in the validation dataset. The dotted line at 45 degrees indicates perfect calibration. **(C)** Receiver operating characteristic curve of the predictive model for good outcome in the validation dataset shows area under the curve of 0.78 [95%CI: 0.66, 0.88].
